# Sex differences in plasma, adipose tissue, and central accumulation of cannabinoids, and behavioural effects of oral cannabis consumption in male and female C57BL/6 mice

**DOI:** 10.1101/2023.05.10.540248

**Authors:** Nada Sallam, Colleen Peterson, Samantha L. Baglot, Yuta Kohro, Tuan Trang, Matthew N. Hill, Stephanie L. Borgland

## Abstract

**Background:** Cannabis edibles are an increasingly popular form of cannabis consumption. Oral consumption of cannabis has distinct physiological and behavioural effects than injection or inhalation. An animal model is needed to understand the pharmacokinetics and physiological effects of oral cannabis consumption in rodents as a model for human cannabis edible use.

**Methods:** Adult male and female C57BL/6 mice received a single dose of commercially available cannabis oil (5 mg/kg THC) by oral gavage. At 0.5-, 1-, 2-, 3-, and 6-hours post-exposure, plasma, hippocampus, and adipose tissue was collected for THC, 11-OH-THC, and THC-COOH measures.

**Results:** We report delayed time to peak THC and 11-OH-THC concentrations in plasma, brain and adipose tissue, which is consistent with human pharmacokinetics studies. We also found sex differences in the cannabis tetrad: (1) female mice had a delayed hypothermic effect 6 hours post-consumption, which was not present in males; (2) females had stronger catalepsy than males; (3) males were less mobile following cannabis exposure, whereas female mice showed no difference in locomotion but an anxiogenic effect at 3h post exposure; and (4) male mice displayed a longer lasting antinociceptive effect of oral cannabis.

**Conclusions:** Oral cannabis consumption is a translationally relevant form of administration that produces similar physiological effects as injection or vaping administration and thus should be considered as a viable approach for examining the physiological effects of cannabis moving forward. Furthermore, given the strong sex differences in metabolism of oral cannabis, these factors should be carefully considered when designing animal studies on the effects of cannabis.

**Significance statement:** Oral delivery of cannabis oil in mice is a translational model that increases plasma, hippocampal, and adipose cannabinoids. Furthermore, oral cannabis and produces lasting psychoactive effects including sex dependent effects on hypothermia, cataplexy, locomotor activity and nociception.

## Introduction

Cannabis edibles are a popular mode of consumption among cannabis users: In 2021, oral cannabis consumption was second only to smoking as the most popular method among those who consumed cannabis(Statistics Canada, 2021). With legalization of cannabis in Canada, self-reported edible cannabis use rose from 32% in 2017 to 53% in 2021(Statistics Canada, 2021). Nearly one third of Canadian and US 16-19 year olds report eating or drinking cannabis in the past 30 days in 2017, 2018, and 2019 (Wadsworth et al., 2022). Furthermore, oral administration is the most common method of use for medicinal users (Sznitman, 2017; Boehnke et al., 2019). Therefore, fully characterized mouse models of oral use are important to understand the acute and long-term physiological consequences of oral cannabis use.

Human pharmacokinetics studies of oral cannabis use have found significant differences in the peak blood and oral fluid concentrations compared to inhalation routes of administration (Swortwood et al., 2017; Spindle et al., 2021; Bidwell et al., 2022). Specifically, the active metabolites ΔL-tetrahydrocannabinol (THC), 11-OH-THC, and THC-COOH peak blood levels are considerably delayed compared to inhalation (Ohlsson et al., 1980; Spindle et al., 2021). This is consistent with reports of subjective “drug effect” peaking 1-2 hours after oral consumption (Vandrey et al., 2017; Schlienz et al., 2020), compared to less than an hour following cannabis inhalation (Ohlsson et al., 1980). Orally administered THC undergoes first-pass metabolism to 11-OH-THC, a metabolite with similar bioavailability at cannabinoid receptor 1 (CB1R) as THC (Nadulski et al., 2005b). Thus, the behavioural and physiological consequences may be distinct from cannabis inhalation or injection. This may also account for differences in some subjective effects of cannabis taken orally versus inhalation (Hart et al., 2002; Spindle et al., 2021).

Preclinical rodent models are useful for studying the effects of cannabis as they allow for fine control of the environment, cannabinoid concentrations, and timing. Moreover, animal research allows us to investigate the biological underpinnings of the effects of cannabis and takes advantage of novel genetic tools and transgenic lines to target neuronal populations, receptors, or enzymes. Many preclinical studies have examined injections of cannabis extract, THC, or CB1R receptor agonists. However, humans rarely administer cannabis through injection, and the pharmacokinetic properties differ between oral and injected cannabis products. Furthermore, while inhalation models have been recently validated (Manwell et al., 2014; Nguyen et al., 2016) and the pharmacokinetics assessed in rats (Baglot et al., 2021), oral cannabis has different pharmacokinetic properties that may have unique behavioural and biological implications. Vaporized cannabis administration models are also complex, challenging to implement, and not accessible to all labs. Therefore, to explore additional ways to model human use, we characterized oral administration in mice as this approach is easily tractable across labs and is still relevant for modelling the effects of cannabis in humans.

Several rodent cannabis studies have used oral consumption, though doses vary considerably between studies (Fairbairn and Pickens, 1979; Abel et al., 1980; Steffens et al., 2005; Mitchell et al., 2021), and it is unclear how these doses compare to human use, given potential differences in rodent metabolism. A 2017 study in Wistar rats examined pharmacokinetic differences in 10 mg/kg THC by route, including oral consumption (Hložek et al., 2017); however, these experiments were only performed in male rats, which did not allow for identification of potential sex differences in the metabolism, tissue disposition, and behavioural response to cannabis. Another study examined the effects of voluntary THC administration in gelatin (1-2 mg/15 ml gelatin) 3 days per week over 33 days and found that the average daily consumption of THC by male and female adolescent rats ranged from 1 to 5 mg/kg and rats consumed greater quantities of the control gelatin than the THC gelatin (Kruse et al., 2019). THC oral administration triggers hypothermia, analgesia and increases in locomotor activity measured after 12 days of chronic use, 1 hour after daily access to THC gelatin (Kruse et al., 2019). Peak plasma concentration following oral administration of 20 mg THC to male subjects, equivalent to 0.33 mg/kg (using 60 kg bodyweight), was 15 ± 10 ng/ml (Grotenhermen, 2003). We converted this to a mouse dose of 4.1 mg/kg using the allometric scaling approach by (Nair and Jacob, 2016), to advise us on the approximate dose to test. Furthermore, an oral THC dose of 5 mg/kg in rats produced locomotor effects and significant antinociception on a tail flick assay (Moore and Weerts, 2022), and produced hypothermia and hypolocomotor activity in male and female C57BL/6 mice (Smoker et al., 2019), and was calculated as the ED_50_ for THC’s antinociceptive effect in male mice (Chesher et al., 1973). However, it is unknown what the tissue distribution or physiological effects oral cannabis are after a single administration in male and female mice. Therefore, for this study, we administered an oral dose of 5 mg/kg whole cannabis THC.

Given that mode of cannabis consumption can produce vastly different pharmacokinetic profiles (Swortwood et al., 2017; Spindle et al., 2021; Bidwell et al., 2022), and oral consumption of cannabis is an increasingly popular method of cannabis consumption among recreational and medical cannabis users, it is critically important to understand the pharmacokinetics and behaviour effect of edible cannabis in mouse models. Here, we analysed the time- and sex-dependent changes in relevant behaviours measuring cannabis intoxication and levels of THC, 11-OH-THC and THC-COOH, in plasma, adipose, and brain tissue in C57BL/6 mice, a common inbred mouse strain, following oral consumption of commercially available cannabis oil containing 95% THC.

## Methods

### Animals

All protocols were in accordance with the ethical guidelines established by the Canadian Council for Animal Care and were approved by the University of Calgary Animal Care Committee. Cannabis use for research was approved under license LIC-IYGANQJY09-2022. Adult (PD90+) male and female C57Bl/6 mice were obtained from Charles River Laboratories (St. Constant, QC, Canada). Mice were group housed (3-4 mice/cage) in a reverse light-dark cycle room (12 h light-dark cycle; lights on: 10 PM MST; lights off: 10 AM MST). All experiments were performed during the dark cycle. Following 2 weeks acclimation, mice were split in to two groups (cannabis and vehicle). For all experiments, commercially available cannabis oil purchased from Tweed Inc. (Smith Falls, Ontario, Canada) containing 25 mg/mL THC in medium chain triglyceride (MCT) oil and less than 1 mg/mL CBD was used. A single dose of 5 mg/kg THC or MCT oil (from coconut oil; Kirkland) was administered by oral gavage at 2.5 hours after lights-off (12:30 PM MST), following 2 hours of fasting to ensure gastric emptying. Mice were habituated to oral gavage, transfer between procedure rooms, and use of the rectal thermometer (for body temperature experiments only) for 3 days prior to test day. Behavioral experiments were performed either immediately after cannabis administration or at the indicated timepoints. Mice were divided into 3 cohorts. One cohort was used to measure rectal temperature, then underwent catalepsy testing and were immediately sacrificed for tissue collection thereafter. Two measurements per mouse were taken (at baseline and another time point). The second cohort was used for the open field test. The third cohort was used for the hot plate test. Different mice were used at different timepoints allowing for collection of plasma and hippocampal samples at each timepoint or to avoid timepoints that would be influenced by prior activity at earlier timepoints (eg hot plate test). Mice were euthanized by CO_2_ directly after behavioural testing. The timepoints were selected based on a previous THC pharmacokinetic study in mice (Torrens et al., 2020; Dumbraveanu et al., 2023).

### Open field

Mice were habituated to the behavioural room for 15 min prior to testing. The open field test was used to assess cannabis-induced changes in locomotion and thigmotaxis. Mice were placed in the center of an arena measuring 30 cm x 30 cm x 30 cm (L x H x W) and allowed to move freely for 10 minutes 0.5-, 1-, 2-, 3-, and 6-hours post cannabis exposure. Movement was recorded with a Basler GenICam mounted above the arena and was analysed using Noldus Ethovision XT 11.5 software (Noldus Information Technology, Leesburg, VA). Locomotor activity was measured as total distance travelled and mean velocity, and the time spent in the center of the chamber. Open field arenas were cleaned with the disinfectant, Virkon between time points and animals to reduce scents.

### Hot plate

Pre-exposure and 3- or 6-hours post-exposure, mice were placed in a 10 cm wide glass cylinder on a hot plate set to 52°C. The latency to reaction (paw licking or jumping), to a maximum of 30 s, was recorded by an observer blinded to the experimental treatment. The apparatus was cleaned with ethanol between animals to remove animal scents.

### Body temperature

Body temperature was taken via a rectal thermometer at baseline and prior to euthanasia for plasma and hippocampus collection at the 0.5-, 1-, 2-, 3-, and 6-hour time points.

### Catalepsy

Catalepsy was assessed by placing the forepaws of the mouse on a bar in the centre of a cage and its hind paws on the floor. The time to move forepaws on or off the bar was recorded as latency to move. The maximum cut off time was 5 min (300 seconds) (Metna-Laurent et al., 2017). The catalepsy apparatus consisted of a modified home cage measuring 28 cm x 12.5 cm x 17.8 cm (L x H x W) with a bar fixed 3.5 cm off the floor in the center of the cage. Catalepsy measurements were taken in the same mice as body temperature measurements, and immediately prior to euthanasia for sample collection at the 0.5-, 1-, 2-, 3-, and 6-hour time points. The catalepsy chamber was cleaned with 20% ethanol between mice.

### Sample preparation

On the day of experiments, mice were fasted for 2 hours to ensure gastric emptying and their baseline temperature and catalepsy assessed. Male (60) and female (60) mice were dosed with commercially available cannabis oil or MCT alone by oral gavage. At either 0.5-, 1-, 2-, 3-, or 6-hours (5-6 mice per time point, per treatment, per sex), rectal body temperature and catalepsy (latency to move) were recorded, the mouse sacrificed, whole blood collected in heparin-lined tubes, the brain rapidly removed, and the hippocampus dissected out. Visceral gonadal adipose tissue was also obtained. The hippocampus or adipose tissue was placed immediately in a tube on dry ice until moved to −80 °C freezer to be stored until analysis. Whole blood was stored on ice for no longer than 2h, then centrifuged at 4 °C for 20 mins at 2500 RPM, the plasma removed and stored at −80 °C until analysis. Samples were prepared as previously described (Baglot et al., 2021). Briefly, samples were thawed at room temperature (plasma) or weighed frozen (hippocampus or adipose tissue) and placed into glass tubes with 2 mL acetonitrile and 100 μL deuterated internal standard (Cerilliant, Round Rock, TX, USA). Hippocampal or adipose samples were first homogenised with a glass rod. Plasma, adipose and brain samples were sonicated in an ice bath for 30 minutes, then stored at −20 °C overnight. Tubes were centrifuged at 1800 RPM at 4 °C for 3-4 minutes, and the supernatant was transferred to a new tube. The supernatant was then evaporated under nitrogen gas, washed with acetonitrile, evaporated and washed again to collect lipids on the tube walls. Samples were then suspended in 1:1 methanol and deionized water, centrifuged twice at 15000 RPM at 4 °C for 20 minutes, and the supernatant stored at −80 C until analysis.

### LC-MS/MS analysis

Samples were prepared for liquid-chromatography-mass spectrometry (LC-MS/MS, multiple reaction monitoring (MRM)) by the SAMs facility at the University of Calgary for measurement of THC, 11-OH-THC, and THC-COOH levels. LC-MS/MS analysis was performed using an Eksigent Micro LC200 coupled to an AB Sciex QTRAP 5500 mass spectrometry (AB Sciex, Ontario, Canada). Analyte concentration (in pmol/µL) were normalized to sample volume/weight and converted to ng/mL.

### Statistics

All data are expressed as mean ± SEM and were analysed and graphed using GraphPad Prism 9.3.1. Normality was determined using a Shapiro-Wilk test. Time-dependent changes in THC, 11-OH-THC, and THC-COOH in males and females were examined using mixed-effects analysis. 3-way ANOVA was used to compare male and female responses of cannabis and time on temperature, catalepsy, and locomotor activity. Time course measurements were not repeated in mice due to either sample collection (THC and metabolite levels), repeated stress (administering rectal probe for temperature) or the effect of repeated measures on behavioural performance (catalepsy, locomotor activity). Hot plate data were compared between baseline and 3 hours or baseline and 6 hours using a repeated measures two-way ANOVA. Sidak’s posthoc tests were used to assess for multiple comparisons. In the figures, asterisks refer to significant p values as follows: *p<0.05, **p<0.01, ***p<0.001, ****p<0.0001.

## Results

### THC, 11-OH-THC, and THC-COOH plasma, brain, and adipose tissue levels

Plasma analyte levels were compared between sexes and examined over time within sexes. A main effect of sex was present in plasma levels of THC (F_(1, 49)_ = 6.134 at p=0.0168, Fig. 1A) with females showing increased levels at 1 hour (p = 0.01) after exposure, and 11-OH-THC (F_(1, 49)_ = 60.81, p<0.0001, Fig. 1B) with significant differences at 1 (p = 0.0002), 2 (p<0.0001), and 3 hours (p = 0.003) after exposure, but not THC-COOH (F_(1, 51)_ = 0.15, p=0.699; Fig. 1C). There was no significant main effect of time on THC levels (F_(4, 49)_ = 2.09, p=0.09). However, there was a significant effect of time on 11-OH-THC levels (F_(4, 49)_ = 3.46, p=0.01) and THC-COOH levels (F_(4, 51)_ = 4.1, p=0.006). In females, T_max_ of plasma THC occurred at 1 hour (C_max_: 27.0 ± 9.6 ng/ml, n = 7), whereas T_max_ of 11-OH-THC (C_max_: 24.3 ± 5.1 ng/ml, n = 6) and peak THC-COOH (C_max_: 25.8 ± 4.3 ng/ml, n = 6) occurred at 2 hours. In males, plasma THC reached T_max_ at 3 hours (C_max_: 12.6 ± 3.1 ng/ml, n = 6), but T_max_ of 11-OH-THC was earlier at 1 hour (C_max_: 1.9 ± 0.9 ng/ml, n = 6) and THC-COOH at 2 hours (C_max_: 22.4 ± 2.2 ng/ml, n = 6). Taken together, females have higher plasma THC and 11-OH THC than males after oral administration.

**Figure 1.**
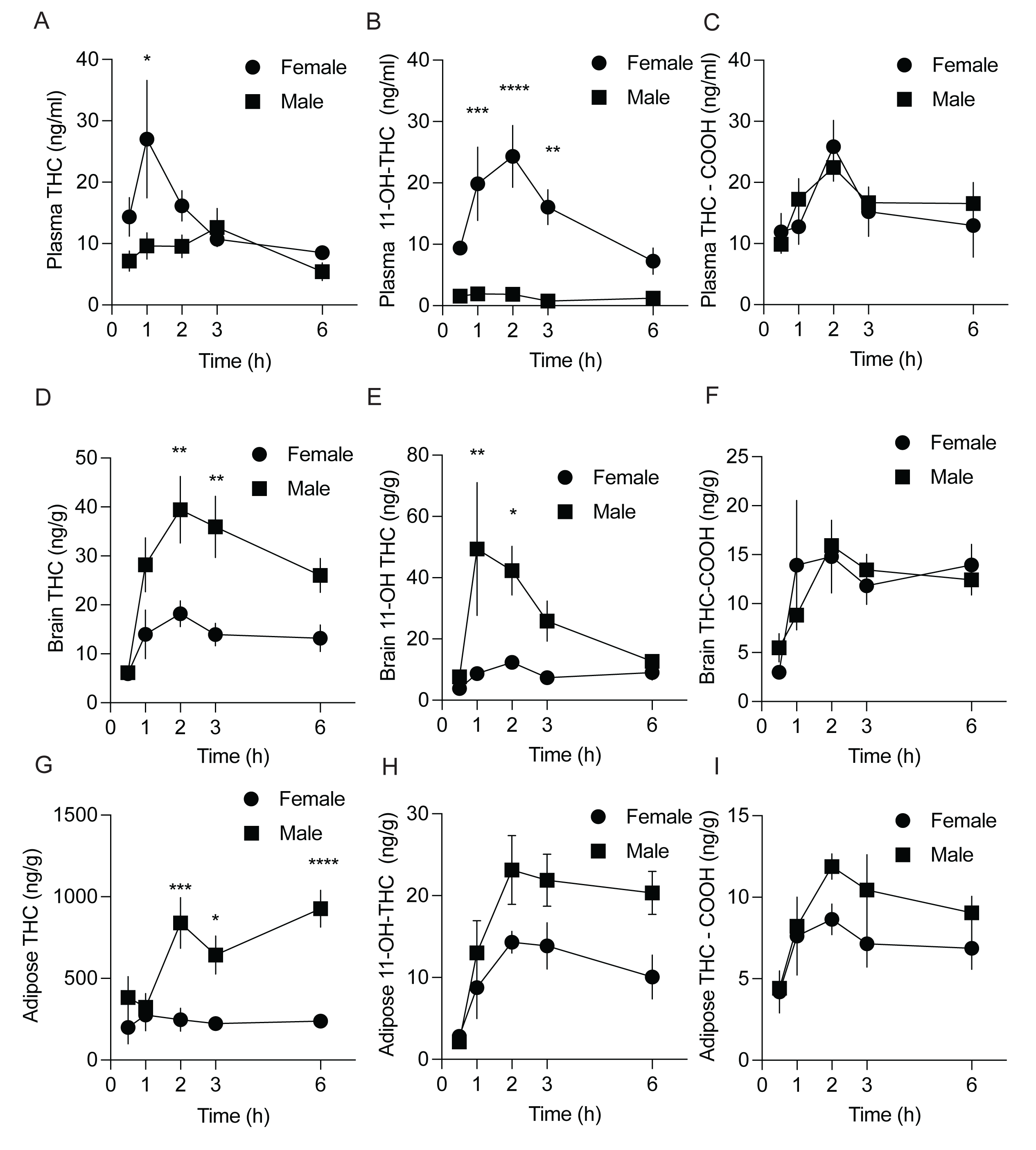
THC,11-OH-THC, THC-COOH concentration in plasma, brain, and adipose tissue. Comparison of male and female plasma levels of (A) THC, (B) 11-OH-THC, and (C) THC-COOH. Comparison of male and female brain (hippocampus) levels of (D) THC, (E) 11-OH-THC, and (F) THC-COOH. Comparison of male and female adipose tissue (G) THC, (H) 11-OH-THC, and (I) THC-COOH. Data are presented as mean +/-SEM, n=5-6.

We next analyzed cannabinoid levels in the brain. We chose to analyze the hippocampus as it is the site for many of the neurobehavioral effects of cannabinoids, has a high expression of cannabinoid receptors (Herkenham et al., 1991). Additionally, hippocampus dissection is straightforward and consistent. In the hippocampus, there was a main effect of sex on THC levels (F_(1,50)_=26.86, p<0.0001; Fig. 1D) with males showing increased levels at 2 (p = 0.003) and 3 hours (p= 0.008) after exposure and 11-OH-THC (F_(1,51)_=15.64 at p = 0.0002; Fig. 1E) with significant differences at 1 (p = 0.003) and 2 hours (p = 0.04) after exposure, but not THC-COOH (F_(1, 50)_ = 0.025 at p=0.88; Fig. 1F). Hippocampal levels of THC reached T_max_ at 2 hours in males (C_max_: 39.4 ± 6.8 ng/g, n = 7) and females (C_max_: 18.2 ± 2.6 ng/g, n = 6). However, hippocampal levels of 11-OH-THC reached T_max_ at 2 hours in females (C_max_: 12.4 ± 1.9 ng/g, n = 6) and 1 hour in males (49.4 ± 21.7 ng/g, n = 6). Hippocampal THC-COOH levels reached T_max_ at 2 hours in males (C_max_: 15.9 ± 1.2 ng/g, n = 7) and females (C_max_: 14.8 ± 3.7 ng/g, n = 6). Taken together, hippocampal levels of THC and 11-OH-THC are higher in males than females, suggesting that uptake to the brain from plasma is much greater in males than females.

In adipose tissue, there was a significant sex x time interaction on THC levels (F_(4,48)_=3.69, p = 0.011; Fig. 1G) and main effects of sex (F_(1,48)_=36.39, p < 0.0001) and time (F_(4,48)_=3.35, p = 0.013), with males showing increased adipose THC accumulation at 2 (p = 0.0005), 3 (p = 0.021) and 6 hours (p < 0.0001). Adipose levels of THC reached T_max_ at 1 hour in females (C_max_: 275.5 ± 70 ng/g, n = 6) and 6 hours in males (C_max_: 927.3 ± 113 ng/g, n = 7). Metabolite levels were considerably lower, with 11-OH-THC reaching T_max_ at 2 hours in females (C_max_: 14.3 ± 1.3 ng/g, n = 6) and males (C_max_: 23.1 ± 4.2 ng/g, n = 6). Similarly, THC-COOH reached T_max_ at 2 hours in females (C_max_: 8.6 ± 0.9 ng/g, n = 6) and males (C_max_: 11.9 ± 0.8 ng/g, n = 6).

#### Cannabis tetrad test battery

We next assessed for sex differences in the physiological and behavioural effects of oral cannabis consumption between male and female mice on 4 measures, body temperature, cataplexy, locomotor activity and nociception. THC administered by injection or inhalation typically induces hypothermia in rats (Nguyen et al., 2020; Ruiz et al., 2021). A 3-way ANOVA indicated there was a significant time x sex interaction (F_(5, 97)_ = 2.72, p = 0.02), a time x cannabis interaction (F_(5, 104)_ = 3.93, p=0.003), but no sex x cannabis interaction (F_(1, 97)_ = 2.67, p = 0.11). There were main effects of time (F_(5, 104)_ = 2.51, p = 0.02), sex (F_(1, 97)_ = 5.97, p = 0.02), and cannabis (F_(1, 104)_ = 5.16, p = 0.03). While there were no significant changes in body temperature after cannabis exposure in males, female mice had lower temperature after cannabis only at the 6-hour timepoint (p = 0.003) (Fig. 2).

**Figure 2.**
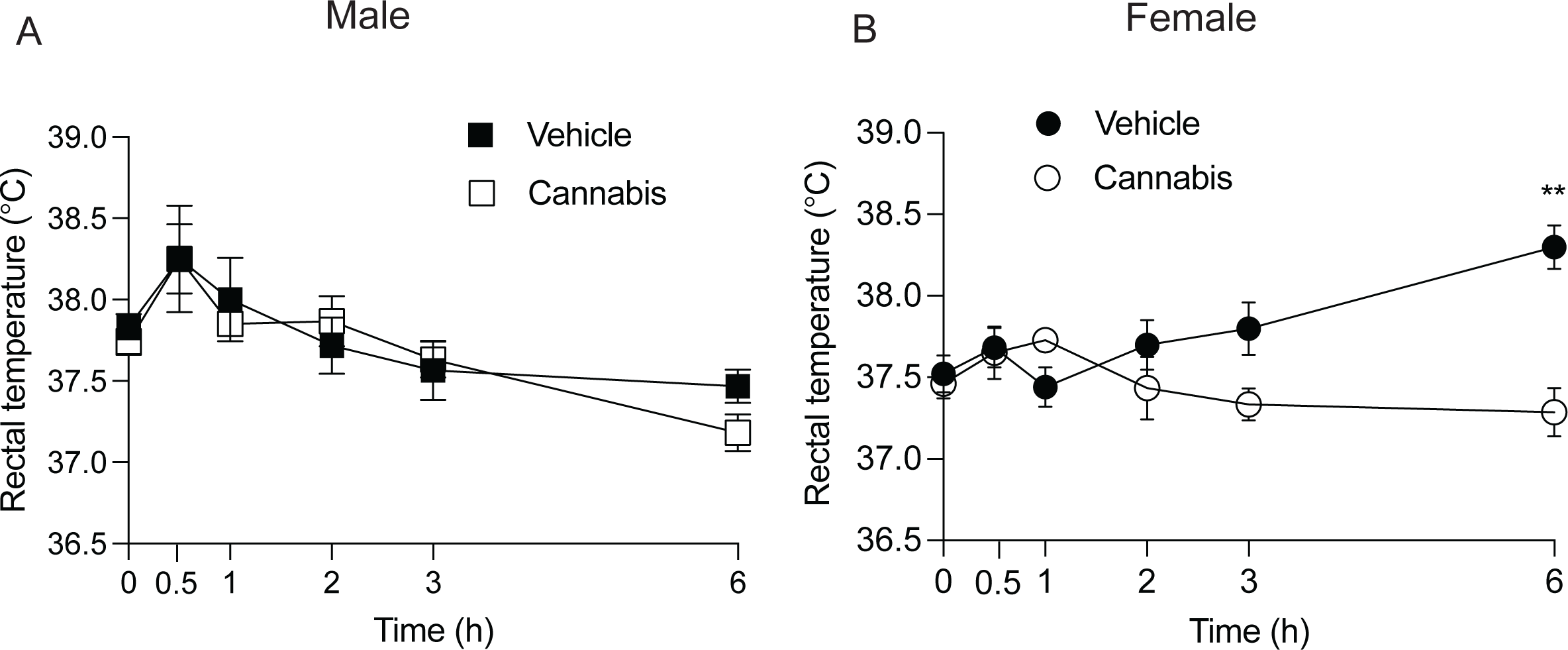
Body temperature following oral cannabis consumption. (A) Male mice do not show a temperature effect of oral cannabis (B) Female mice show a delayed hypothermic effect of oral cannabis 6 hours post-consumption. Data are presented as mean +/-SE, n=6-7.

To determine if oral cannabis exposure influenced catalepsy in male or female mice, we placed mice on a horizontal bar and measured the time until forepaws were off the bar. A 3-way ANOVA indicated that there were significant time x cannabis (F_(5, 117)_ = 13.10, p < 0.0001), sex x cannabis (F_(1, 90)_ = 39.34, p < 0.0001), and time x sex x cannabis interactions (F_(5, 90)_ = 10.56, p < 0.0001). There were also main effects of time (F_(5, 117)_ = 11.38, p < 0.0001), sex (F_(1, 90)_ = 28.98, p < 0.0001), and cannabis (F_(1, 117)_ = 22.29, p< 0.0001). Oral consumption of cannabis oil resulted in significant catalepsy in male (Figure 3A) and female mice (Figure 3B). However, the cataleptic effect of cannabis was greater, had an earlier onset and was longer lasting in females with significant differences from vehicle at 2 (p < 0.0001) and 3 hours (p < 0.0001) after oral administration.

**Figure 3.**
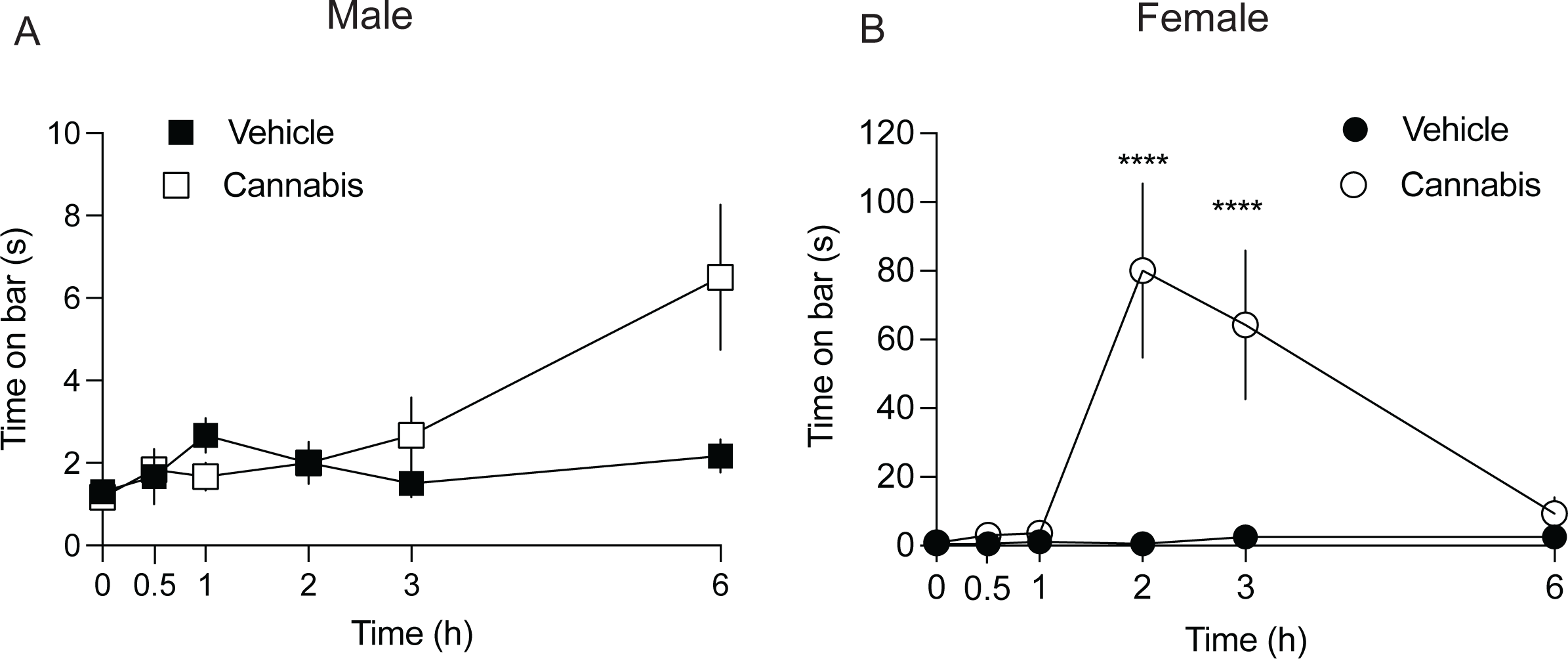
Catalepsy following oral cannabis consumption. (A) Male mice have a significantly increased latency to move after placement on the catalepsy bar at 6 hours post-cannabis consumption. (B) In female mice, the catalepsy is greatest at two hours and only persists to 3 hours post-cannabis consumption. Data are presented as mean +/-SEM, n=6-7.

To determine if oral cannabis exposure altered locomotor behaviour, we measured distance travelled within 10 min on the open field test. A 3-way ANOVA indicated that there were significant main effects of time (F_(4,179)_ = 23.47, p < 0.0001), cannabis (F_(1,170)_ = 10.71, p = 0.0013), and sex (F_(1,170)_ = 19.74, p < 0.0001) on distance travelled, but no significant interactions of time x cannabis (F_(4,170)_ = 1.08, p = 0.37), time x sex (F_(4,170)_ = 0.24, p = 0.92), cannabis x sex (F_(1,170)_ = 0.77, p = 0.38). Whole cannabis oil significantly decreased locomotion in the open field task at 3 hours in male (Fig.4A, p = 0.041), but not female mice (Fig. 4B, p = 0.19).

**Figure 4.**
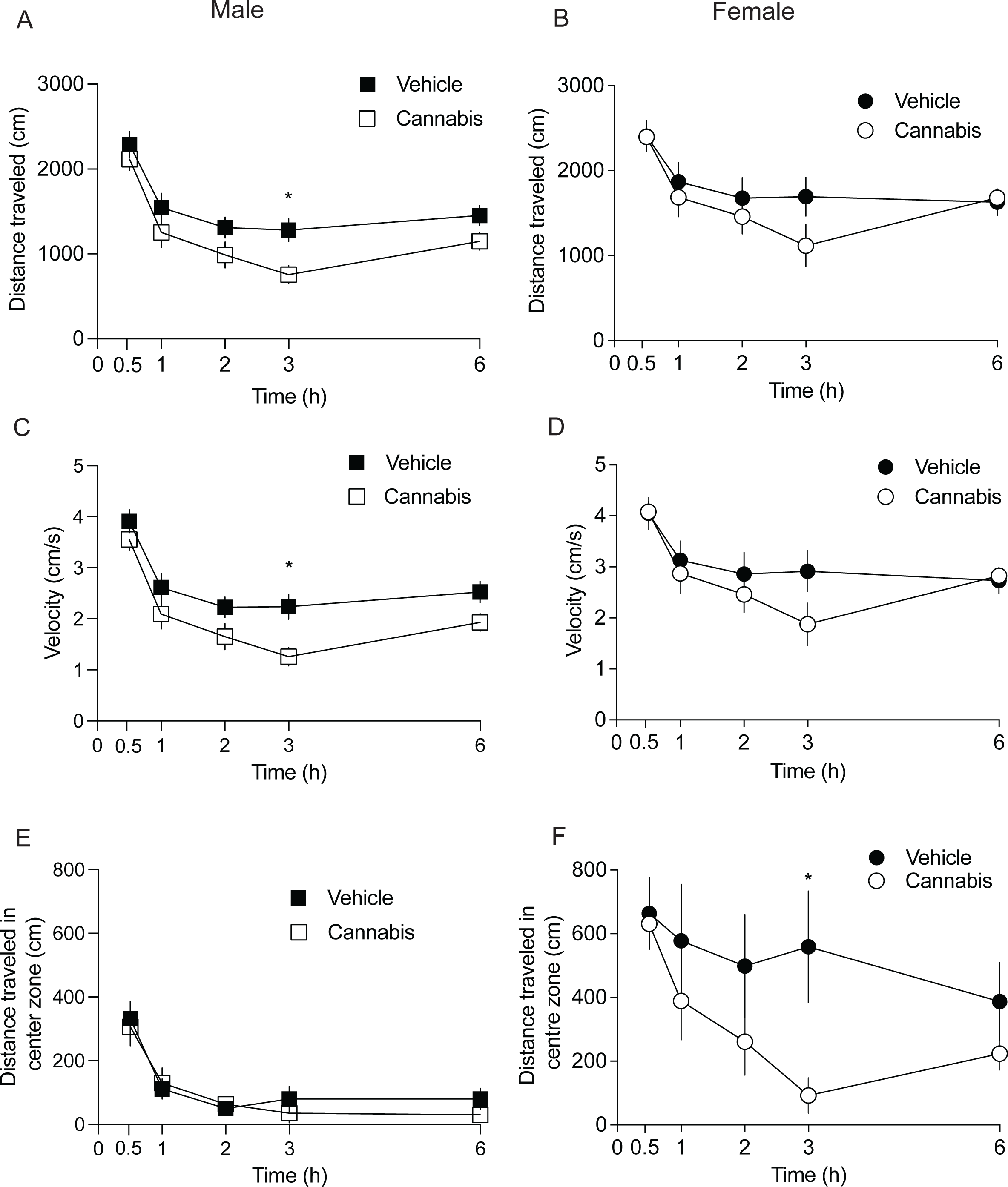
Locomotor effects of oral cannabis consumption. (A) Male mice show a hypolocomotor effect of oral cannabis consumption at 3 hours. (B) Female offspring did not show a significant locomotor effect of cannabis. Velocity was not significantly different in male (C) or female (D) mice. There was no effect of cannabis on distance travelled in the centre of the open field apparatus for male mice (E), but (F) female mice had decreased distance traveled in the centre of the open field apparatus 3 hours post-cannabis consumption. Data are presented as mean +/-SEM, n=9-10 mice.

We then examined the effects of oral cannabis consumption on velocity in male (Fig. 4C) and female mice (Fig. 4D). A 3-way ANOVA indicated that there were significant main effects of time (F_(4,170)_ = 23.18, p = 0.38), cannabis (F_(1,170)_ = 12.03, p = 0.0007), and sex (F_(1,170)_ = 19.18, p < 0.0001). However, there were no interactions of time x cannabis (F_(4,170)_ = 1.24, p =0.29), time x sex (F_(4,170)_ = 0.26, p =0.91), or cannabis x sex (F_(1,170)_ = 1.21, p =0.27). Oral cannabis significantly decreased velocity at 3 hours in male (Fig.4C, p = 0.03), but not female mice (Fig. 4D, p = 0.36).

We next measured distance travelled in the center zone which is suggestive of anxiolytic behaviour as mice become more thigmotaxic, remaining near the walls of the open field apparatus with higher anxiety. A 3-way ANOVA indicated that there were significant main effects of time (F_(4,126)_ = 6.35, p = 0.0001), cannabis (F_(1,126)_ = 6.84, p = 0.01), and sex (F_(1,126)_ = 46.44, p < 0.0001) and a significant cannabis x sex interaction (F_(1,126)_ = 4.92, p = 0.028). There were no interactions of time x cannabis (F_(4,126)_ = 0.73, p =0.57) or time x sex (F_(4,126)_ = 0.20, p = 0.93). Oral cannabis significantly decreased time in center at 3h in female (Fig.4F, p = 0.046), but not male mice (Fig. 4E, p = 0.91). This suggests that oral cannabis may be anxiogenic in female mice.

##### Anti-nociception (hot plate test)

We tested for cannabis-induced anti-nociception using a hot plate test. Oral cannabis consumption produced a sex-specific anti-nociceptive effect. At 3 hours post-exposure in male mice (Fig. 5A), there was a cannabis x time interaction (F_(1, 17)_ = 5.47, p = 0.032), but no main effects of cannabis F_(1, 17)_ = 1.6, p = 0.22) or time (F_(1, 17)_ = 1.22, p = 0.16). A Sidak’s posthoc test revealed a significant increase in latency to evoke nociceptive behaviors (paw licking or jumping) of cannabis at 3 hours (p = 0.036). In female mice (Fig. 5B), there was a cannabis x time interaction (F_(1, 16)_ = 4.5, p = 0.049), a main effect of time (F_(1, 17)_ = 6.89, p = 0.018), but no main effect of cannabis (F_(1, 16)_ = 1.35, p = 0.26). A Sidak’s posthoc test revealed a significant increase in latency to evoke nociceptive behaviors of cannabis at 3 hours (p = 0.008). At 6 hours post cannabis exposure, male mice displayed significantly increased latency to evoke nociceptive behaviors (cannabis x time interaction (F_(1, 13)_ = 11.63, p = 0.005); cannabis effect: F_(1, 13)_ = 7.27, p = 0.02), time effect: F_(1, 13)_ = 10.02, p = 0.007); Sidak’s posthoc: p = 0.002), whereas female mice showed no difference (cannabis x time interaction (F_(1, 18)_ = 0.001, p = 0.97); cannabis effect: F_(1, 18)_ = 0.29, p = 0.59), time effect: F_(1, 18)_ = 0.31, p = 0.58), suggesting that the antinociceptive effects of oral cannabis last longer in males than females.

**Figure 5.**
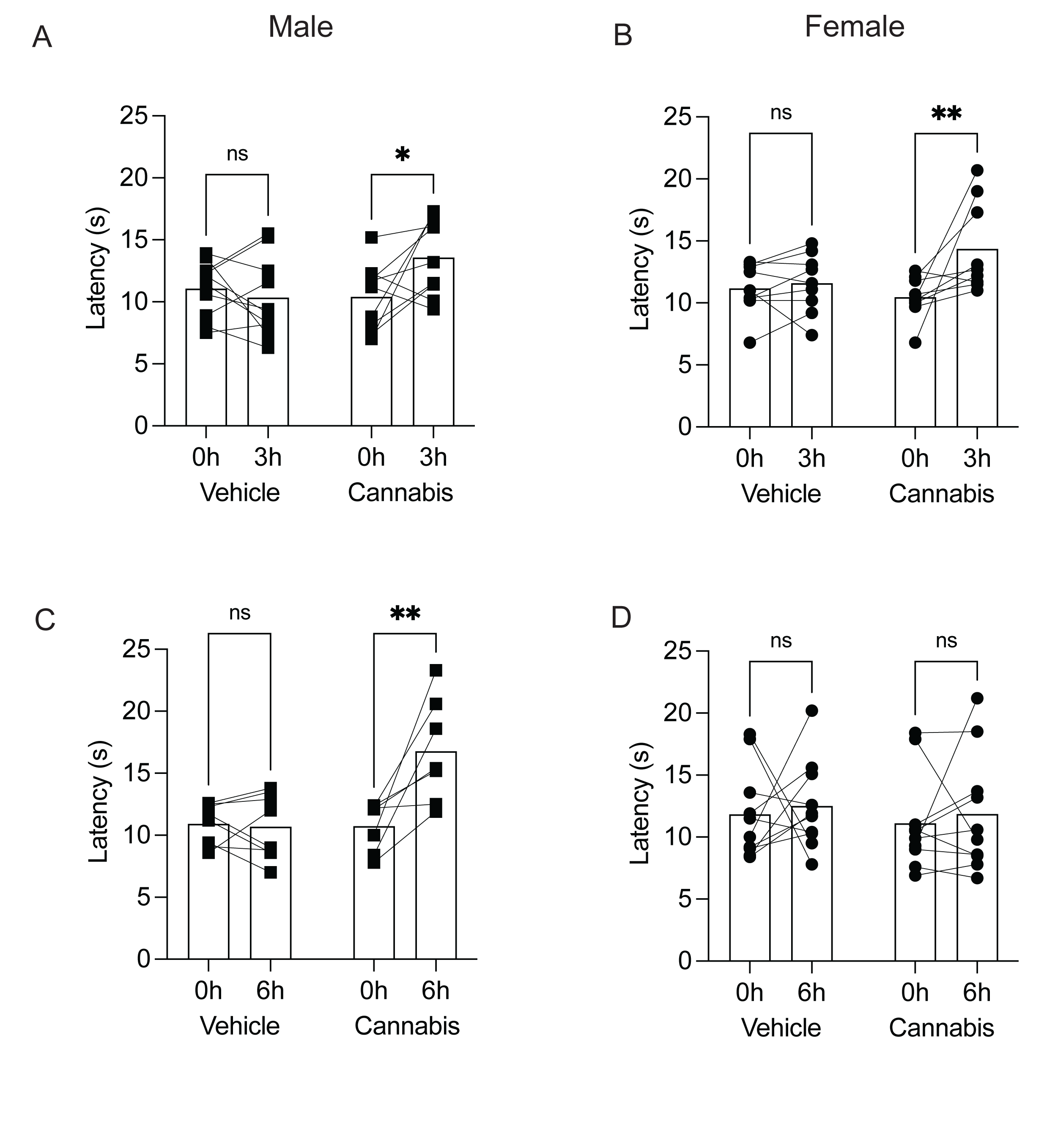
Nociceptive effects of oral cannabis consumption. Male (A) or female (B) mice have increased latency to evoke nociceptive behaviours (paw licking or jumping) on the hot plate test at 3h post consumption. At 6h post consumption, male mice have increased latency to paw licking on the hot plate test (C), but this effect was absent in female mice (D). Bars represent mean and symbols represent individual responses before or after vehicle (shaded bars) or cannabis (open bars) consumption. Data are presented as mean +/-SEM, n=7-10 mice.

## Discussion

In the present study we demonstrated that oral cannabis produces significant sex differences in plasma and brain concentrations of THC, 11-OH-THC, and THC-COOH. Furthermore, these differences in pharmacokinetic effect of oral cannabis were observed in behavioural responses, such that females had a stronger hypothermic, cataleptic, and anxiogenic response than males, but a shorter lasting antinociceptive effect. Thus, oral cannabis shows significant sex differences in the levels and tissue dispersal of THC and its metabolite 11-OH-THC, as well as behavioural effects in the cannabis tetrad.

### Plasma and brain distribution of THC and metabolite concentrations

We have examined the pharmacokinetics of oral consumption of whole cannabis oil (5 mg/kg THC) in male and female C57BL/6 mice in plasma and brain tissue. We found that plasma THC levels reached T_max_ earlier and reached higher C_max_ in female mice compared to male mice. Notably, the psychoactive metabolite 11-OH-THC was also had a significantly greater C_max_ in female than male mice, but THC-COOH plasma levels were similar in male and female mice. Higher plasma THC concentration in females than males is consistent with injection and inhalation studies (Nguyen et al., 2020; Baglot et al., 2021; Ruiz et al., 2021). However, while plasma THC concentration rapidly increases after vapour administration in rats (peak within 15-35 min), injection studies had a slower rise in concentration with peaks occurring between 35-90 min (Nguyen et al., 2020; Baglot et al., 2021; Ruiz et al., 2021). Notably, oral administration resulted in a delayed peak concentration occurring at 1 hour for females and 2 hours for males. This time course is similar to oral administration of 20 mg/kg THC in male C57BL6 mice (Dumbraveanu et al., 2023), 15 mg/kg THC from a full spectrum cannabis extract in male and female C57BL/6 mice (Anderson et al., 2021), and 10 mg/kg THC in male Wistar rats (Hložek et al., 2017). In contrast, mice have faster conversion to THC-COOH compared to Wistar rats (Hložek et al., 2017), with peak levels around 2 hours in brain and plasma as opposed to 4 hours in rat plasma and 8 hours in rat brain. This may be due to differences in mouse and rat metabolism of THC, or due to the lower dose used in our study. The metabolite 11-OH-THC was significantly lower in concentration than THC plasma levels, consistent with that observed in male rats (Hložek et al., 2017) and mice (Dumbraveanu et al., 2023). Furthermore, there was a significant sex difference in plasma 11-OH-THC with greater levels in females compared to males. This is consistent with that observed with vapor or injection administration in rodents (Tseng et al., 2004; Baglot et al., 2021; Ruiz et al., 2021) and oral or vapour administration in humans (Sholler et al., 2021). There may exist sex differences in THC metabolism which could explain the increased female plasma 11-OH-THC levels. Liver microsomes harvested from female adult Sprague Dawley rats preferentially oxidised THC to 11-OH-THC, whereas male rat liver microsomes produce a broader array of metabolites (Narimatsu et al., 1991, 1992). While this, if occurring in C57Bl/6 mice, may explain the increased levels of 11-OH-THC in plasma of female mice, it does not account for the increase of THC and 11-OH-THC in male over female brain.

In men, a single oral dose of 20 mg THC resulted in a C_max_ of 14 ± 9.7 ng/ml (Wall et al., 1983) or 12.4 ± 3.4 ng/ml (Schwilke et al., 2009), which is similar to that observed in male mice after an oral dose of 5 mg/kg (C_max_ = 12.6 ± 3.1 ng/ml). Few studies have examined sex differences in the pharmacokinetics of human oral cannabis consumption. Thus, it is difficult to compare the similarity of response in female mice to that in women. However, one study found a C_max_ of 9.4 ± 4.5 ng/ml in women given an oral dose of 15 mg THC (Wall et al., 1983). Our study found a twofold increase in plasma THC over male mice with a C_max_ of 27 ± 9.6 ng/ml.

We have also shown significant sex differences in the peak hippocampal and adipose tissue concentration of THC and time course of THC and 11-OH-THC. While peak plasma THC was lower and peaked later in males than in females, hippocampal and adipose tissue concentrations of THC and 11-OH-THC were higher in males than female animals. This might indicate that THC is more rapidly absorbed into male mouse brain and adipose tissue from plasma compared to females. One potential explanation is that there are differences in blood-brain-barrier levels of transporters which can bind THC and its metabolites resulting in differences in cannabinoid efflux. Specifically, the transporters ABCB1a/b and ABCG2 are implicated in brain disposition of THC, such that a knockout results in significantly higher levels of THC in the brain (Spiro et al., 2012). It is unclear whether these transporters also transport 11-OH-THC. Consistent with previous work (Kreuz and Axelrod, 1973; Torrens et al., 2020), we found that accumulation of THC in adipose tissue of male mice was 10x greater than that in brain. Increased accumulation of lipophilic THC in adipose tissue in male mice may be due to greater intra-abdominal retroperitoneal and gonadal fat stores in male compared to female mice (Shi et al., 2007).

The data herein are also generally consistent with human pharmacokinetic studies that have measured plasma or whole blood THC. Oral consumption of cannabis results in peak blood or plasma levels of THC and 11-OH-THC 1-3 hours post-consumption (Poyatos et al., 2020). However, many human pharmacokinetic studies combine data from males and females, and thus it is unclear if there are strong sex differences in the metabolism of cannabis after oral consumption in humans as observed in rodents. One study found significantly higher plasma THC and 11-OH-THC levels in females and shorter time to peak for each than males after a single 10 mg THC edible dose with or without CBD, which persisted after controlling for weight (Nadulski et al., 2005a).

### Sex differences in cannabis tetrad

We also saw significant differences in the behavioural and physiological effects of oral cannabis consumption in the cannabis tetrad test. Specifically, we saw that female mice had a delayed hypothermic effect of cannabis 6 hours post-consumption, which was not present in males; that while both male and female mice displayed a cataleptic effect of cannabis, it was more pronounced in females; males had slightly decreased activity following cannabis exposure, whereas female mice showed no difference in locomotion, although a decrease in distance traveled in the center zone was evident. Finally, while both male and female mice displayed an antinociceptive effect of oral cannabis, it was longer lasting in males.

Despite lower brain THC and 11-OH-THC levels, female mice seem to be more sensitive to the cataleptic and hypothermic effects of THC and 11-OH-THC. The lack of hypothermic effect of cannabis in males is not unprecedented. A recent publication which examined the cannabis tetrad response after oral cannabis in rats found that at a 5.6 mg/kg THC dose, male rats showed no hypothermic effect of cannabis at any point, but a significant hypothermic effect in females after 5 hours (Moore and Weerts, 2022), similar to what we have reported here. While this study also noted that CBD increased body temperature in male rats, the doses used in this study (3, 10, and 30 mg/kg CBD) are considerably higher than the CBD present in the whole cannabis oil we have used, which contained <1 mg/mL CBD. Thus, CBD does not likely contribute to the sex differences in the hypothermic effect of cannabis we report here. Cataleptic effects of cannabinioids have been reported from intraperitoneal injections (Tseng and Craft, 2001; Tseng et al., 2004) or oral administration of THC (Cohn et al., 1972), intraperitoneal injection of 11-OH −THC or the CBR1/2 agonist CP55940 (Tseng and Craft, 2001) in rats. Consistent with our results, there was a greater cataleptic effect of intraperitoneal THC in female rats compared to male rats (Tseng et al., 2004). However, when the cytochrome P450 inhibitor blocked conversion of THC to 11-OH-THC, this sex difference was abolished, suggesting that this sex difference in catalepsy is likely due to the psychoactive metabolite. Notably, we observed significantly lower levels of brain 11-OH-THC in females than males, therefore factors other than metabolite concentration likely also play a role in cataleptic response. One factor for this difference may include the contribution of the estrous cycle, which has been previously shown to impact cannabinoid receptor expression in some regions (de Fonseca et al., 1994; Castelli et al., 2014), but which we did not control for.

Some behavioural sex differences are consistent with our observation of increased male brain levels of THC and its primary active metabolite, 11-OH-THC. Namely, male mice reduced their locomotor activity, and had longer lasting antinociception following oral cannabis consumption. Decreased locomotor activity was also observed in male Wistar rats 2 hours after oral THC (Tseng et al., 2004). Furthermore, consistent with our results, a recent study of oral cannabis tetrad effects in rats also found that male rats had a stronger anti-nociceptive effect of 5.6 mg/kg THC oral cannabis in response to thermal stimuli 5 hours post-consumption (Moore and Weerts, 2022); however, both sexes showed a similar anti-nociceptive response at 2 hours post-consumption and in response to a physical stimulus.

### Limitations and conclusions

Previous pharmacokinetic studies have largely used rats, making it difficult to directly compare pharmacokinetic responses of cannabis as species differences between rats and mice likely contribute to differences in metabolism and distribution of cannabis. Furthermore, our mice were fasted prior to oral gavage to ensure gastric emptying to control for food consumption prior to cannabis dosing. In humans, oral consumption of cannabis while fasted results in earlier peak THC and 11-OH-THC times (Lunn et al., 2019); thus, the time to peak THC and metabolites we report may be shorter than expected compared to fed mice. Furthermore, although we administered cannabis oil via oral gavage to tightly control the time that mice received cannabis, we recognise this is a stressful procedure (Brown et al., 2000). Therefore, our behavioural results may differ from those produced by a voluntary oral cannabis consumption model. This study only used a single dose of THC. There is evidence of biphasic effects of THC on behaviour with injection models (Patel and Hillard, 2006; Rubino et al., 2007), however this has not been well-investigated in oral models. Future studies should address the possibility of biphasic effects with different THC concentrations in the oral administration model. Finally, because this study design collected both blood and brain samples at each time point rather than taking repeated measures from the same animal over time, we were unable to effectively measure repeated pharmacokinetic parameters such as the half-life of elimination, volume of distribution, clearance, or elimination rate constant. Future studies should address these parameters with oral administration in mice in comparison with other routes of administration as well as other developmental periods.

In conclusion, using a commercially available cannabis oil at a common dose used in rodent studies (5 mg/kg THC), we have characterised the pharmacokinetics and behavioural effects of oral cannabis consumption in male and female C57BL/6 mice. We have also demonstrated significant sex differences in the pharmacokinetics of THC, and report different behavioural responses to oral cannabis consumption in male versus female mice in agreement with previous research (Smoker et al., 2019; Wiley et al., 2021). It is unclear to what extent sex differences in response to cannabis are due to differences in THC metabolism versus the impact of THC and its metabolites on sex differences in neural circuits, and/or expression of cannabinoid receptors within these circuits, underlying these behaviours. These findings should be considered by researchers interested in using preclinical models of cannabis consumption as this oral administration model is low cost, easy to use, and translationally relevant. We encourage future animal and human oral cannabis studies to carefully consider potential sex differences in their experimental design and analyses given the significant sex-dependent effects observed here.

## Acknowledgements

The authors would like to acknowledge the Southern Alberta Mass Spectrometry facility. This research was performed at the University of Calgary which is located on the unceded traditional territories of the people of the Treaty 7 region in Southern Alberta, which includes the Blackfoot Confederacy (including the Siksika, Piikuni, Kainai First Nations), the Tsuut’ina, and the Stoney Nakoda (including the Chiniki, Bearspaw, and Goodstoney First Nations). The City of Calgary is also home to Metis Nation of Alberta, Region III.

## Funding

This work is supported by a University of Calgary VPR Catalyst Grant and a Matheson Centre Research Grant on Cannabis (S.L.B), Alberta Innovates Research Grant - mCannabis and CIHR (PJT-162271) to T.T., CIHR Catalyst grant-Cannabis to M.H., and Tier 1 Canada Research Chair (S.L.B. 950-232211). C.P. was supported by an Alberta Innovates Graduate Studentship in Health Innovation.

## Conflict of Interest Statement

The authors declare no competing financial or other conflicts of interest.

## Data Availability Statement

Data is available upon request.

## Notes

### Competing Interest Statement

The authors have declared no competing interest.

### Summary of Updates

Some references are updated

## References

Abel EL, Dintcheff BA, Day N (1980) Effects of marihuana on pregnant rats and their offspring. Psychopharmacology 71:71–74.

Anderson LL, Etchart MG, Bahceci D, Golembiewski TA, Arnold JC (2021) Cannabis constituents interact at the drug efflux pump BCRP to markedly increase plasma cannabidiolic acid concentrations. Sci Rep 11:14948.

Baglot SL, Hume C, Petrie GN, Aukema RJ, Lightfoot SHM, Grace LM, Zhou R, Parker L, Rho JM, Borgland SL, McLaughlin RJ, Brechenmacher L, Hill MN (2021) Pharmacokinetics and central accumulation of delta-9-tetrahydrocannabinol (THC) and its bioactive metabolites are influenced by route of administration and sex in rats. Sci Rep 11:23990.

Bidwell LC, Karoly HC, Torres MO, Master A, Bryan AD, Hutchison KE (2022) A naturalistic study of orally administered vs. inhaled legal market cannabis: cannabinoids exposure, intoxication, and impairment. Psychopharmacology 239:385–397.

Boehnke KF, Scott JR, Litinas E, Sisley S, Clauw DJ, Goesling J, Williams DA (2019) Cannabis Use Preferences and Decision-making Among a Cross-sectional Cohort of Medical Cannabis Patients with Chronic Pain. The journal of pain 20:1362–1372.

Brown AP, Dinger N, Levine BS (2000) Stress produced by gavage administration in the rat. Contemporary topics in laboratory animal science 39:17–21.

Castelli MP, Fadda P, Casu A, Spano MS, Casti A, Fratta W, Fattore L (2014) Male and female rats differ in brain cannabinoid CB1 receptor density and function and in behavioural traits predisposing to drug addiction: effect of ovarian hormones. Curr Pharm Des 20:2100–2113.

Chesher GB, Dahl CJ, Everingham M, Jackson DM, Marchant-Williams H, Starmer GA (1973) The effect of cannabinoids on intestinal motility and their antinociceptive effect in mice. British Journal of Pharmacology 49:588–594.

Cohn RA, Barratt E, Pirch JH (1972) Differences in behavioral responses of male and female rats to marijuana. Proceedings of the Society for Experimental Biology and Medicine Society for Experimental Biology and Medicine (New York, NY) 140:1136–1139.

de Fonseca FR, Cebeira M, Ramos JA, Martín M, Fernández-Ruiz JJ (1994) Cannabinoid receptors in rat brain areas: Sexual differences, fluctuations during estrous cycle and changes after gonadectomy and sex steroid replacement. Life Sciences 54:159–170.

Dumbraveanu C, Strommer K, Wonnemann M, Choconta JL, Neumann A, Kress M, Kalpachidou T, Kummer KK (2023) Pharmacokinetics of Orally Applied Cannabinoids and Medical Marijuana Extracts in Mouse Nervous Tissue and Plasma: Relevance for Pain Treatment. Pharmaceutics 15.

Fairbairn JW, Pickens JT (1979) The oral activity of delta’-tetrahydrocannabinol and its dependence on prostaglandin E2. British journal of pharmacology 67:379–385.

Grotenhermen F (2003) Pharmacokinetics and pharmacodynamics of cannabinoids. Clin Pharmacokinet 42:327–360.

Hart CL, Ward AS, Haney M, Comer SD, Foltin RW, Fischman MW (2002) Comparison of smoked marijuana and oral Δ9-tetrahydrocannabinol in humans. Psychopharmacology 164:407–415.

Herkenham M, Lynn AB, Johnson MR, Melvin LS, Costa B de, Rice KC (1991) Characterization and localization of cannabinoid receptors in rat brain: a quantitative in vitro autoradiographic study. J Neurosci 11:563–583.

Hložek T, Uttl L, Kadeřábek L, Balíková M, Lhotková E, Horsley RR, Nováková P, Šíchová K, Štefková K, Tylš F, Kuchař M, Páleníček T (2017) Pharmacokinetic and behavioural profile of THC, CBD, and THC+CBD combination after pulmonary, oral, and subcutaneous administration in rats and confirmation of conversion in vivo of CBD to THC. European Neuropsychopharmacology 27:1223–1237.

Kreuz DS, Axelrod J (1973) Delta-9-tetrahydrocannabinol: localization in body fat. Science (New York, NY) 179:391–393.

Kruse LC, Cao JK, Viray K, Stella N, Clark JJ (2019) Voluntary oral consumption of Δ9-tetrahydrocannabinol by adolescent rats impairs reward-predictive cue behaviors in adulthood. Neuropsychopharmacology 44:1406–1414.

Lunn S, Diaz P, O’Hearn S, Cahill SP, Blake A, Narine K, Dyck JRB (2019) Human Pharmacokinetic Parameters of Orally Administered Δ(9)-Tetrahydrocannabinol Capsules Are Altered by Fed Versus Fasted Conditions and Sex Differences. Cannabis and cannabinoid research 4:255–264.

Manwell LA, Ford B, Matthews BA, Heipel H, Mallet PE (2014) A vapourized Δ(9)-tetrahydrocannabinol (Δ(9)-THC) delivery system part II: comparison of behavioural effects of pulmonary versus parenteral cannabinoid exposure in rodents. Journal of pharmacological and toxicological methods 70:112–119.

Metna-Laurent M, Mondésir M, Grel A, Vallée M, Piazza P-V (2017) Cannabinoid-Induced Tetrad in Mice. Curr Protoc Neurosci 80:9.59.1-9.59.10.

Mitchell VA, Harley J, Casey SL, Vaughan AC, Winters BL, Vaughan CW (2021) Oral efficacy of Δ(9)-tetrahydrocannabinol and cannabidiol in a mouse neuropathic pain model. Neuropharmacology 189:108529.

Moore CF, Weerts EM (2022) Cannabinoid tetrad effects of oral Δ9-tetrahydrocannabinol (THC) and cannabidiol (CBD) in male and female rats: sex, dose-effects and time course evaluations. Psychopharmacology 239:1397–1408.

Nadulski T, Pragst F, Weinberg G, Roser P, Schnelle M, Fronk E-M, Stadelmann AM (2005a) Randomized, double-blind, placebo-controlled study about the effects of cannabidiol (CBD) on the pharmacokinetics of Delta9-tetrahydrocannabinol (THC) after oral application of THC verses standardized cannabis extract. Therapeutic drug monitoring 27:799–810.

Nadulski T, Sporkert F, Schnelle M, Stadelmann AM, Roser P, Schefter T, Pragst F (2005b) Simultaneous and sensitive analysis of THC, 11-OH-THC, THC-COOH, CBD, and CBN by GC-MS in plasma after oral application of small doses of THC and cannabis extract. Journal of Analytical Toxicology 29:782–789.

Nair AB, Jacob S (2016) A simple practice guide for dose conversion between animals and human. J Basic Clin Pharm 7:27–31.

Narimatsu S, Watanabe K, Matsunaga T, Yamamoto I, Imaoka S, Funae Y, Yoshimura H (1992) Cytochrome P-450 isozymes involved in the oxidative metabolism of delta 9-tetrahydrocannabinol by liver microsomes of adult female rats. Drug metabolism and disposition: the biological fate of chemicals 20:79–83.

Narimatsu S, Watanabe K, Yamamoto I, Yoshimura H (1991) Sex difference in the oxidative metabolism of Δ9-tetrahydrocannabinol in the rat. Biochemical Pharmacology 41:1187–1194.

Nguyen JD, Aarde SM, Vandewater SA, Grant Y, Stouffer DG, Parsons LH, Cole M, Taffe MA (2016) Inhaled delivery of Δ(9)-tetrahydrocannabinol (THC) to rats by e-cigarette vapor technology. Neuropharmacology 109:112–120.

Nguyen JD, Creehan KM, Grant Y, Vandewater SA, Kerr TM, Taffe MA (2020) Explication of CB(1) receptor contributions to the hypothermic effects of Δ(9)-tetrahydrocannabinol (THC) when delivered by vapor inhalation or parenteral injection in rats. Drug and alcohol dependence 214:108166.

Ohlsson A, Lindgren J-E, Wahlen A, Agurell S, Hollister LE, Gillespie HK (1980) Plasma delta-9-tetrahydrocannabinol concentrations and clinical effects after oral and intravenous administration and smoking. Clinical Pharmacology & Therapeutics 28:409–416.

Patel S, Hillard CJ (2006) Pharmacological evaluation of cannabinoid receptor ligands in a mouse model of anxiety: further evidence for an anxiolytic role for endogenous cannabinoid signaling. J Pharmacol Exp Ther 318:304–311.

Poyatos L, Pérez-Acevedo AP, Papaseit E, Pérez-Mañá C, Martin S, Hladun O, Siles A, Torrens M, Busardo FP, Farré M (2020) Oral Administration of Cannabis and Δ-9-tetrahydrocannabinol (THC) Preparations: A Systematic Review. Medicina (Kaunas, Lithuania) 56:309.

Rubino T, Sala M, Viganò D, Braida D, Castiglioni C, Limonta V, Guidali C, Realini N, Parolaro D (2007) Cellular Mechanisms Underlying the Anxiolytic Effect of Low Doses of Peripheral Δ9-Tetrahydrocannabinol in Rats. Neuropsychopharmacol 32:2036–2045.

Ruiz CM, Torrens A, Lallai V, Castillo E, Manca L, Martinez MX, Justeson DN, Fowler CD, Piomelli D, Mahler S V (2021) Pharmacokinetic and pharmacodynamic properties of aerosolized (“vaped”) THC in adolescent male and female rats. Psychopharmacology 238:3595–3605.

Schlienz NJ, Spindle TR, Cone EJ, Herrmann ES, Bigelow GE, Mitchell JM, Flegel R, LoDico C, Vandrey R (2020) Pharmacodynamic dose effects of oral cannabis ingestion in healthy adults who infrequently use cannabis. Drug and Alcohol Dependence 211:107969.

Schwilke EW, Schwope DM, Karschner EL, Lowe RH, Darwin WD, Kelly DL, Goodwin RS, Gorelick DA, Huestis MA (2009) Δ9-Tetrahydrocannabinol (THC), 11-Hydroxy-THC, and 11-Nor-9-carboxy-THC Plasma Pharmacokinetics during and after Continuous High-Dose Oral THC. Clin Chem 55:2180–2189.

Shi H, Strader AD, Woods SC, Seeley RJ (2007) The effect of fat removal on glucose tolerance is depot specific in male and female mice. American journal of physiology Endocrinology and metabolism 293:E1012–20.

Sholler DJ, Strickland JC, Spindle TR, Weerts EM, Vandrey R (2021) Sex differences in the acute effects of oral and vaporized cannabis among healthy adults. Addict Biol 26:e12968.

Smoker MP, Mackie K, Lapish CC, Boehm SL (2019) Self-administration of edible Δ9-tetrahydrocannabinol and associated behavioral effects in mice. Drug Alcohol Depend 199:106–115.

Spindle TR, Martin EL, Grabenauer M, Woodward T, Milburn MA, Vandrey R (2021) Assessment of cognitive and psychomotor impairment, subjective effects, and blood THC concentrations following acute administration of oral and vaporized cannabis. Journal of Psychopharmacology 35:786–803.

Spiro AS, Wong A, Boucher AA, Arnold JC (2012) Enhanced brain disposition and effects of Δ9-tetrahydrocannabinol in P-glycoprotein and breast cancer resistance protein knockout mice. PloS one 7:e35937–e35937.

Statistics Canada (2021) Canadian Cannabis Survey 2021: Summary.

Steffens S, Veillard NR, Arnaud C, Pelli G, Burger F, Staub C, Zimmer A, Frossard J-L, Mach F (2005) Low dose oral cannabinoid therapy reduces progression of atherosclerosis in mice. Nature 434:782–786.

Swortwood MJ, Newmeyer MN, Andersson M, Abulseoud OA, Scheidweiler KB, Huestis MA (2017) Cannabinoid disposition in oral fluid after controlled smoked, vaporized, and oral cannabis administration. Drug Testing and Analysis 9:905– 915.

Sznitman SR (2017) Do recreational cannabis users, unlicensed and licensed medical cannabis users form distinct groups? International Journal of Drug Policy 42:15– 21.

Torrens A, Vozella V, Huff H, McNeil B, Ahmed F, Ghidini A, Mahler SV, Huestis MA, Das A, Piomelli D (2020) Comparative Pharmacokinetics of Δ9-Tetrahydrocannabinol in Adolescent and Adult Male Mice. J Pharmacol Exp Ther 374:151–160.

Tseng AH, Craft RM (2001) Sex differences in antinociceptive and motoric effects of cannabinoids. European journal of pharmacology 430:41–47.

Tseng AH, Harding JW, Craft RM (2004) Pharmacokinetic factors in sex differences in Delta 9-tetrahydrocannabinol-induced behavioral effects in rats. Behavioural brain research 154:77–83.

Vandrey R, Herrmann ES, Mitchell JM, Bigelow GE, Flegel R, LoDico C, Cone EJ (2017) Pharmacokinetic Profile of Oral Cannabis in Humans: Blood and Oral Fluid Disposition and Relation to Pharmacodynamic Outcomes. Journal of Analytical Toxicology 41:83–99.

Wadsworth E, Craft S, Calder R, Hammond D (2022) Prevalence and use of cannabis products and routes of administration among youth and young adults in Canada and the United States: A systematic review. Addict Behav 129:107258.

Wall ME, Sadler BM, Brine D, Taylor H, Perez-Reyes M (1983) Metabolism, disposition, and kinetics of delta-9-tetrahydrocannabinol in men and women. Clinical Pharmacology & Therapeutics 34:352–363.

Wiley JL, Barrus DG, Farquhar CE, Lefever TW, Gamage TF (2021) Sex, species and age: Effects of rodent demographics on the pharmacology of Δ9-tetrahydrocanabinol. Prog Neuropsychopharmacol Biol Psychiatry 106:110064.

